# Genomic signatures of honey bee association in an acetic acid symbiont

**DOI:** 10.1101/367490

**Authors:** Eric A. Smith, Irene L. G. Newton

## Abstract

Honey bee queens are central to the success and productivity of their colonies; queens are the only reproductive members of the colony, and therefore queen longevity and fecundity can directly impact overall colony health. Recent declines in the health of the honey bee have startled researchers and lay people alike as honey bees are agriculture’s most important pollinator. Honey bees are important pollinators of many major crops and add billions of dollars annually to the US economy through their services. One factor that may influence queen and colony health is the microbial community. Although honey bee worker guts have a characteristic community of bee-specific microbes, the honey bee queen digestive tracts are colonized by a few bacteria, notably an acetic acid bacterium not seen in worker guts: *Bombella apis.* This bacterium is related to flower-associated microbes such as *Saccharibacter floricola* and other species in the genus *Saccharibacter,* and initial phylogenetic analyses placed it as sister to these environmental bacteria. We used comparative genomics of multiple honey bee-associated strains and the nectar-associated *Saccharibacter* to identify genomic changes associated with the ecological transition to bee association. We identified several genomic differences in the honey bee-associated strains, including a complete CRISPR/Cas system. Many of the changes we note here are predicted to confer upon them the ability to survive in royal jelly and defend themselves against mobile elements, including phages. Our results are a first step towards identifying potential benefits provided by the honey bee queen microbiota to the colony’s matriarch.

## Introduction

The honey bee (*Apis mellifera*) is extremely important economically because of the pollination services it provides to numerous agricultural crops. As a result, there is increasing interest in determining how the microbiome supports and influences bee function. While a honey bee colony is made up of bees with diverse roles, or castes, the majority of studies on bee microbiomes have focused on workers specifically. The microbial community of worker bees consists of eight to ten core bacterial species [1–5]. The characterization of these groups led to speculation about their role in honey bee health and whether or not they provision nutrients [3] or assist in the breakdown of plant-derived carbohydrates [6], as is the case in other insect-microbe interactions [7, 8]. There has also been speculation as to the role of the microbiome in resistance to pathogens, as microbial communities have been shown to protect the bumble bee (*Bombus terristris*) from the parasite *Crithidia bombi* [9]. Honey bee-associated microbes interact with each other in diverse ways both *in vitro* and *in vivo*, suggesting that they may interact syntrophically within workers [2, 10]. While these studies focused on honey bee workers are intriguing, it is surprising that only recently was the microbiome of queen bees throughout development recently characterized [11–13].

Interestingly, the microbial community associated with queen bees is vastly different than associated workers and comprises a large percentage of acetic acid bacteria, a group of bacteria present only at very small percentages in workers. One of the primary bacteria that differentiate queens from workers is the recently described *Bombella apis* [14], formerly *Parasaccharibacter apium* [15]. *Bombella apis* is in the family *Acetobacteraceae* and occupies defined niches within the hive, including: queen guts, nurse hypopharyngeal glands, nurse crops, and royal jelly, and is only rarely found outside of these areas [16–18]. Evidence suggests that it might play a role in protecting developing larvae and worker bees from fungal pathogens [15, 19]. Given that *B. apis* makes up a large proportion of the queen gut microbiome, it is possible that it plays important roles in queen health and physiology, as well [13, 20].

*Bombella apis* is part of a clade of acetic acid bacteria (AAB, a group within the family *Acetobacteraceae*) that contains both free-living and bee-associated members. Comparative genomics, then, can give us insights into the changes associated with the transition to bee-association in this clade. This comparison can also help elucidate what sets *B. apis* apart from closely related species and the role(s) it might be playing in the hive environment. To that end, we used the genomes of eight *B. apis* strains [21–24], a *Bombella sp.* genome assembly, as well as five genomes of the closely-related genus *Saccharibacter* [25, 26] and a genome of the bumblebee symbiont, *Bombella intestini* [27], to begin to tease apart the unique capabilities of *B. apis*. Insights gained here could prove critical in determining the factors responsible for maintaining queen health in colonies and could ultimately lead to the development of interventions to improve queen health and mitigate the detrimental impacts of queen failure on this economically critical species.

## Materials and methods

### *Phylogenetic relationship of* Bombella *and* Saccharibacter

To determine the placement of *Bombella* and *Saccharibacter* strains used here, we initially used the 16S rRNA gene sequences from the Silva database [28–30] that met the following criteria: 1) from a species belonging to either *Bombella* or *Saccharibacter,* 2) of length at least 1200 bases, and 3) sequence quality >90. Additionally, the 16S rRNA gene sequence for *Gluconobacter oxydans* was included as an outgroup. The 16S rRNA sequences for *Bombella intestini* [27] and *Bombella apis* [14] were included. We used BLAST to find the 16S rRNA gene sequences in the *Bombella* and *Saccharibacter* genomes (Table 1) to pull out their respective 16S rRNA sequences for use in this phylogeny. All sequences were aligned using the SINA aligner [31]; parameters used were set using the --auto option. A maximum likelihood phylogeny was constructed using RAxML with the GTRGAMMA substitution model and 1000 bootstrap replicates (v8.2.11, [32]). The final tree was visualized using FigTree (v1.4.2, http://tree.bio.ed.ac.uk/software/figtree/).

**Table 1.**
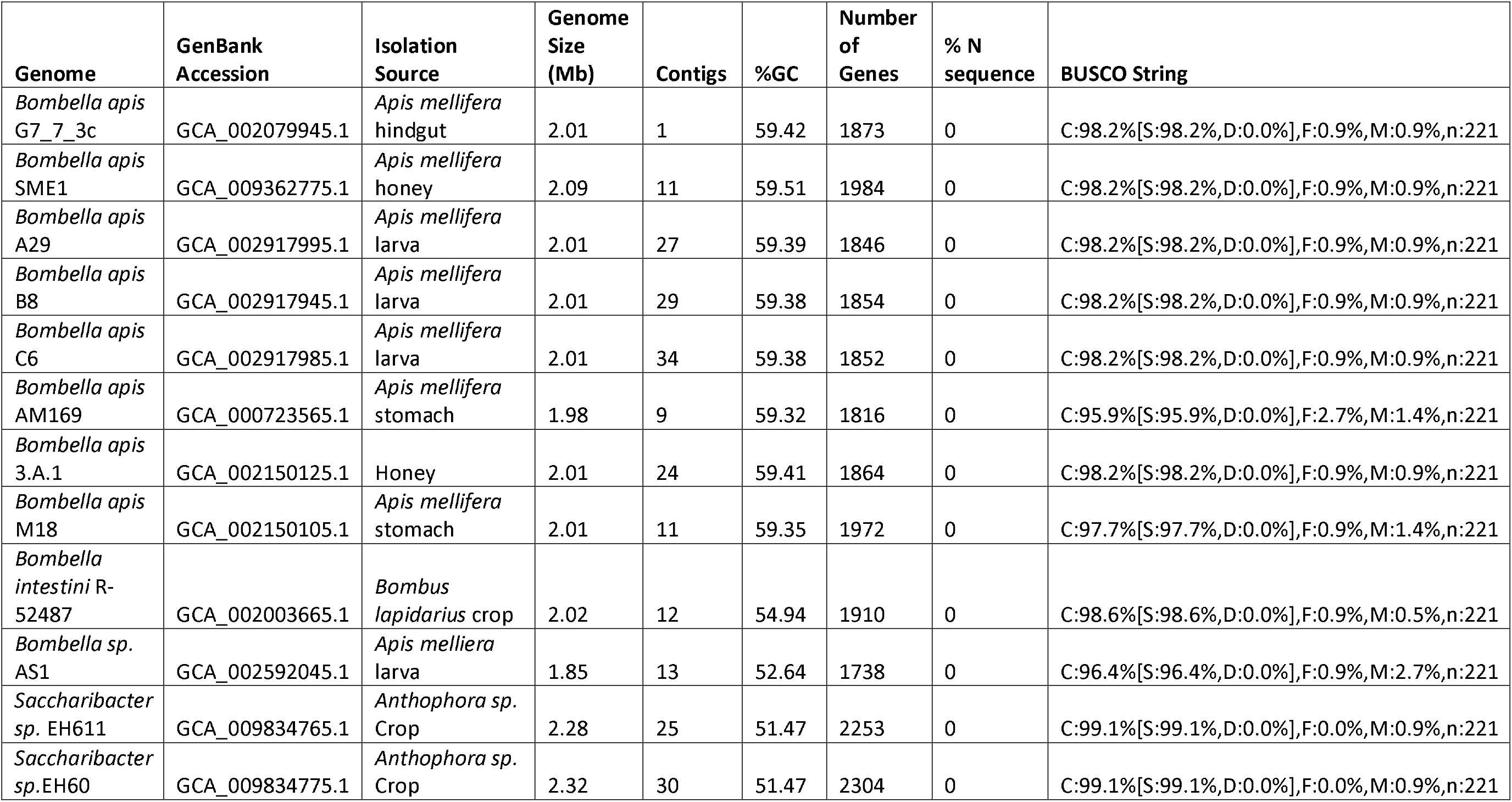
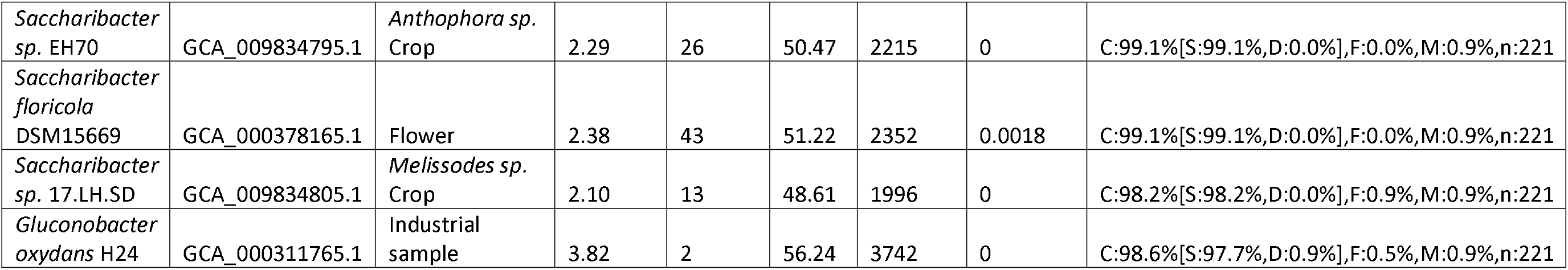
Genome names, accession number, and isolation sources for genomes used in these analyses. *Bombella apis* G7_7_3c is the *Bombella apis* reference genome.

### Orthology analysis

To facilitate downstream analyses, we clustered genes from all genomes in Table 1 – plus *Gluconobacter oxydans* H24 as an outgroup – into groups of orthologous genes (GOGs) using OrthoMCL (v.2.0.9, [33]) (Table S1). Amino acid sequences were downloaded from NCBI and clustering was performed using default OrthoMCL parameters, namely percentMatchCutoff=50 and evalueExponentCutoff=−5. These clusters were then classified as single-copy orthologs (defined as containing exactly one representative from each genome), variable (defined as missing a representative from at least one genome and having varying numbers of representatives from each of the other genomes), multi-copy ortholog (containing at least one representative from each genome, but multiple copies from at least one genome), or genome-specific (containing at least two genes that all came from the same genome) using an in-house Perl script.

### Bombella *and* Saccharibacter *core ortholog phylogeny*

We constructed a phylogeny using concatenated amino acid alignments of all single-copy GOGs. The amino acid sequences were aligned using the MAFFT L-INS-I algorithm (v7.310, [34]), and alignments were then concatenated, and used to construct a maximum likelihood phylogeny using RAxML with substitution model PROTGAMMALGF and 1000 bootstrap replicates (v8.2.11, [32]). The final tree was visualized using FigTree (v1.4.2, http://tree.bio.ed.ac.uk/software/figtree/).

### Calculation of genomic similarity

To determine relatedness and species assignment, we calculated genome-wide Average Nucleotide Identity (gANI) and aligned fraction (AF) for each pairwise comparison using ANIcalculator.[35] Predicted transcript sequences for each pairwise comparison were passed to the software, which output gANI and AF in each direction for the pairwise comparison. As gANI and AF can vary depending on the direction of comparison due to differences in genome length, we report the average of the pairwise calculations in each direction.

### Synteny analysis

Before our analysis of synteny, we subjected all of our assemblies to Quast v. 5.0.2 run with default parameters on contigs from our genomes compared to the circularized genome of *B. apis* strain G7_7_3 using the --pe1 and --pe2 flags to include raw reads and calculate genome completeness as well as determine potential misassemblies. Few potential within-contig misassemblies were identified in each genome, with most having no flagged misassemblies. We identified some within-contig misassemblies for the following strains: 9 for *Bombella apis* A29, and 7 for *B. apis* C6 and *B. apis* SME1. Importantly, we cannot rule out misassembly of the G7_7_3 genome or natural genomic inversions and rearrangements between these strains over time. Also, no misassemblies were identified in regions hosting HGTs mentioned below. We then used Mauve [36, 37] to determine the syntenic regions between the *Bombella* spp genomes. The *Bombella apis* G7_7_3c reference genome is resolved to a single chromosome, so it was used as the reference sequence in Mauve’s “move contigs” tool, and the likely order and orientation of contigs in the other genomes was determined. To facilitate downstream analyses, the output of Mauve’s “move contigs” tool was used to order, orient, and concatenate contigs into single pseudo-chromosomes for each genome. Structural rearrangements were then visualized using Mauve’s built-in graphical interface.

### Annotation of CRISPR arrays and phage sequences

Pseudo-chromosomes for each genome were uploaded to CRISPRFinder to determine location and sequence of CRISPR arrays [38]. To assess whether CRISPR arrarys differ genome-to-genome (an indication that the arrays were incorporated after the strains diverged) we used an in-house Perl script to determine the maximum intergenomic percent identity of spacer sequences by aligning each spacer from a given genome to every spacer in every other genome and calculating percent identity. We also used the SEA-PHAGES databased to identify phages from which these spacers might be derived (Table S6). We used PHAge Search Tool Enhanced Release (PHASTER) [39, 40] to define phage-like regions. Any region determined to be “questionable” or “intact” by PHASTER was labeled as likely to be of phage origin.

### Determination of bee-associated bacteria-specific orthologs

We identified all GOGs that contained at least one gene from each genome of *Bombella* spp. and no genes from *Saccharibacter* spp. We then took the *B. apis* G7_7_3c genome representative for each of these GOGs and got KEGG annotations for as many as possible using BlastKOALA [41]. Any hit that was given a definitive KO number by BlastKOALA was considered valid. For those genes that we were not able to get KEGG annotations, we used NCBI’s BLAST to aid in determining potential function of these bee-associated bacteria-specific genes (Table S2). This list of genes and their potential functions was then manually inspected to hypothesize genes that may have allowed for the transition to bee-association.

### Analysis of horizontal gene transfers

To determine whether or not genes in any of the *Bombella* spp. genomes were horizontally transferred, we employed a combination of sequence-composition, phylogenetic, and synteny approaches. We mapped genes of particular interest (e.g. genes unique to certain clades, species, or strains) to their locations on the linear pseudo-chromosomes constructed during synteny analysis. Additionally, we calculated the %GC for each gene. We then determined how many standard deviations each gene was from its genome-wide mean %GC. The third prong of this analysis involved identifying genes that were phylogenetically aberrant. To do this, we used Darkhorse [42] to calculate the lineage probability index (LPI) for each gene. LPI measures the likelihood that a particular gene was inherited vertically, from the ancestor of the species of interest. Higher LPIs indicate a higher likelihood that the gene is ancestral and has not been horizontally transferred into the resident genome, while lower LPIs indicate that horizontal gene transfer (HGT) may have occurred. Darkhorse calculates LPI using the taxonomy string for the best BLAST hits for each gene. We ran LPI twice, once including BLAST hits to *Bombella* and *Saccharibacter* subject sequences and once excluding such hits from the analysis. In doing so, genes with orthologs in close relatives of the *Bombella*/*Saccharibacter* clade (likely vertically inherited) will have high LPIs in both analyses, whereas genes without orthologs in close relatives of the clade (potentially horizontally transferred) will have a high LPI in the analysis that included *Bombella* and *Saccharibacter* BLAST hits, but a low LPI in the analysis that excluded those BLAST hits. For each gene, then, we can calculate the difference in LPI between the two analyses to determine how far from the genome-wide mean LPI difference (in standard deviations) each gene is. In doing so, genes that are likely to be horizontally transferred will have a larger discrepancy between LPI values than genes that were vertically inherited. We then classified regions as likely to be HGTs if they met the following criteria: 1) a block of at least three syntenic genes that show interesting phylogenetic distributions (e.g. unique to clade, species, or strain) where 2) a majority of genes in the region are at least 1 standard deviation from the mean %GC or LPI difference (or both).

### Domain annotation of genes of interest

We used HHpred (https://toolkit.tuebingen.mpg.de/#/tools/hhpred, [43]) to determine domain architecture and gain an understanding of potential function of the genes in each HGT. For genes of interest that were part of a GOG, all members of the GOG were first aligned using the MAFFT L-INS-I algorithm (v7.310, [34]). These multiple sequence alignments (or single amino acid sequences in the case of strain-unique genes) were then uploaded to HHpred’s online tool and homology was determined using HMMs in the COG_KOG_v1.0, Pfam-A_v31.0, and SMART_v6.0 databases; only domains scoring above 60% probability are discussed here. Gene models for each region of interest were then constructed and visualized using the HHpred results and in-house R scripts to “draw” the gene models. The fifth HGT identified (HGT5) occurs at the junction of two contigs in the linear pseudo-chromosomes we constructed. The abutting ends of each contig have annotations for partial pseudogenes, such that when they are joined a complete gene is created. A BLAST search using the nucleotide sequence of this gene against the NCBI nr database was used to determine a putative function.

## Results

### Saccharibacter *and* Bombella *are sister clades*

To robustly determine the relationship between *Bombella* and *Saccharibacter* spp., we constructed a maximum likelihood phylogeny using 16S rRNA sequences. Our final tree largely agrees with previously published phylogenies for this group (Figure 1) [18, 19]. Sequences were largely grouped into two monophyletic clades by genus. *Bombella* plus *Saccharibacter* comes out as sister to the *Gluconobacter* outgroup. The taxonomic nomenclature of this entire group has recently been revised but clearly *Bombella* and *Saccharibacter* are separate clades [14, 27].

**Fig. 1.**
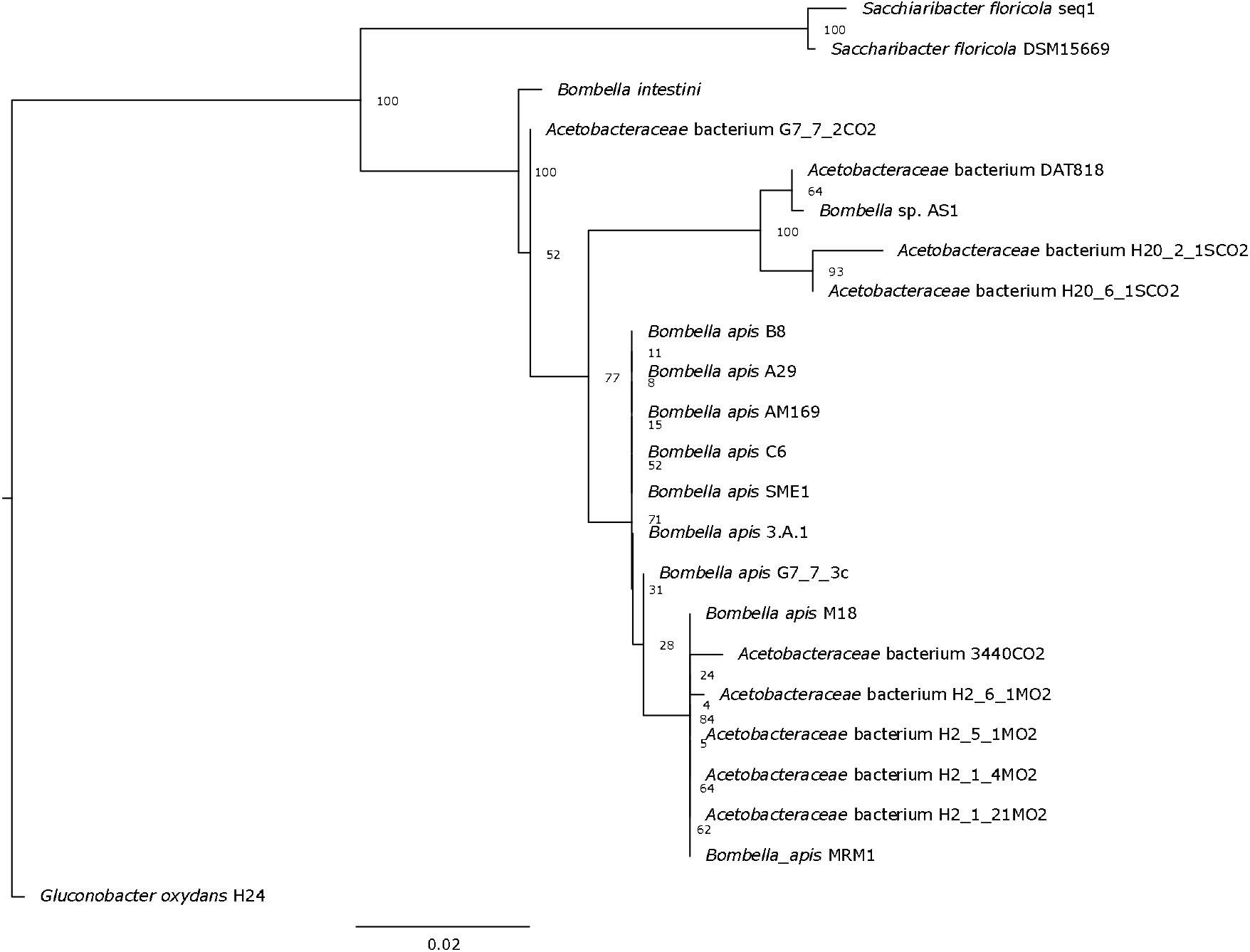
Maximum likelihood phylogenetic tree of *Bombella* and *Saccharibacter* species constructed from full-length 16S rRNA sequences, *Gluconobacter oxydans* as an outgroup. Bootstrap scores are indicated at each node.

### *Core ortholog phylogeny of* Bombella *and* Saccharibacter *strains*

We used OrthoMCL (v2.0.9 [33]) to define groups of orthologous genes (GOGs) using the *Bombella* and *Saccharibacter* genomes listed in Table 1; *Gluconobacter oxydans* H24 was used as an outgroup. In total, 3,017 GOGs were defined, with an average of 10.3 genes per GOG. Of these, 1,209 GOGs were present as single copies in every genome in the analysis, while an additional 34 GOGs were present in every genome, but in varying numbers in each genome. There were 1,526 GOGs that were variable (i.e. missing a representative from at least one genome, and present in varying numbers in the other genomes); and 248 GOGs that consisted of at least two genes that all came from the same genome (Table S1).

To better resolve the phylogenetic relationships between *Bombella* spp. and *Saccharibacter* spp., we constructed a second maximum likelihood phylogeny using aligned and concatenated amino acid sequences of the 1,209 single-copy GOGs (Figure 2). This robustly supported amino acid phylogeny broadly agrees with our previously constructed 16S phylogeny. In the core ortholog tree, *B. intestini* interrupts the monophyly of honey bee-associated *Bombella* genomes. Notably, this tree groups *B. intestini* more closely to the majority of the *Bombella apis* strains, while *P. apium* AS1 is more distantly related, and possibly a different species. Similar to the 16S tree, we again see quite short branch lengths within the honey bee-associated acetic acid bacteria, particularly among those in the clade including all *Bombella apis* genomes (Figure 2).

**Fig. 2.**
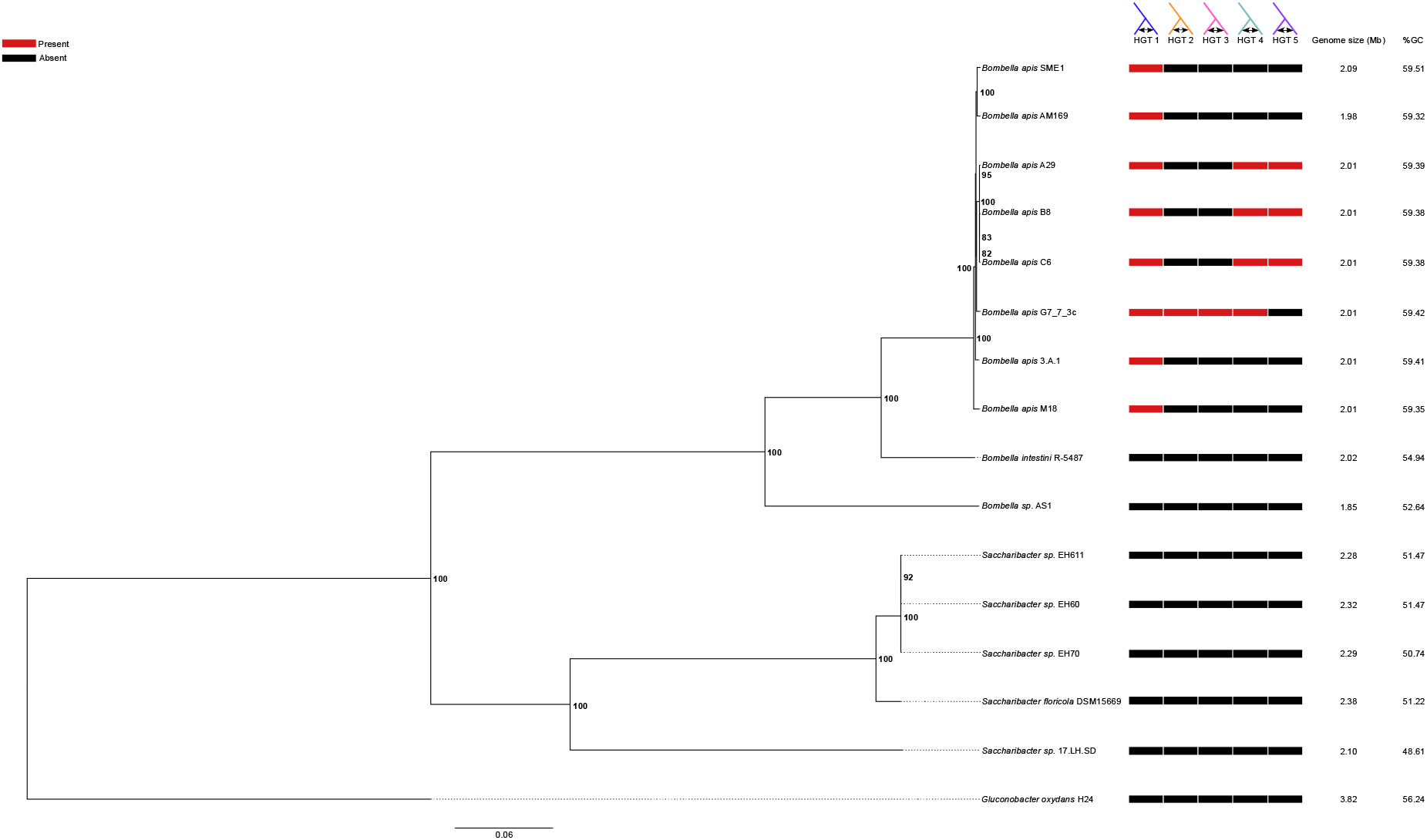
Maximum likelihood phylogenetic tree of the *Bombella*/*Saccharibacter* clades constructed from concatenated amino acid alignment of 1259 single copy orthologous genes. Bootstrap scores are indicated at each node. Colored boxes represent the presence (red) or absence (black) of each of 7 genomic regions of interest. Genome size and %GC are also displayed.

### Identification of sequences of phage origin

Movement and insertion of bacteriophage sequences in a genome can have profound effects on the evolution of that genome [44–46]. Mobile genetic elements can also provide insight into the lifestyle of a bacterium, as the fraction of mobile DNA varies significantly with host ecology [47]. We identified two regions of phage origin among the 15 genomes analyzed. One region in *S. floricola* was identified as “intact”, stretching approximately 60 kilobases (kb), and containing 89 proteins, all of which are identified by BLAST hits as being of phage origin or hypothetical. Synteny alignments indicate that this region is unique to *S. floricola* and not contained in any other genome. Likewise, OrthoMCL did not cluster any of the genes within this region with any other genes in our analysis, further supporting the idea that these genes are unique to *S. floricola*. The second region is in the *B. apis* G7_7_3c genome and was identified as “questionable” (one step below “intact”). This phage region is approximately 31kb long and contains 40 proteins, all of which were identified as either being of phage origin or hypothetical. Like Phage 1, synteny mapping and OrthoMCL clustering showed that this region is unique to *B. apis* G7_7_3c and shows no homology to any of the other genomes (Figure 2).

### *Signatures of honey bee association in the* B. apis *genomes*

To identify genes associated with the transition to honey bee association, we identified GOGs that contained at least one gene from each *Bombella* spp. and were also missing in *Saccharibacter* spp. There were a total of 1,542 GOGs containing at least one gene from each of the aforementioned genomes, but only 74 were also missing in all *Saccharibacter* spp. We determined the putative functions of these genes using the *B. apis* G7_7_3c genome representative for each GOG (Table S2). It should be noted that all annotations discussed from here forward are putative and require further functional characterization.

Several bee-associated unique genes stood out as particularly interesting, the first being gluconolactonase. Lactonases, such as gluconolactonase, reversibly catalyze the hydrolysis of lactones (such as gluconolactone) to the hydroxyl acid form (such as gluconic acid). Gluconolactone is found in both honey and royal jelly and is thought to be partially responsible for the antibacterial properties of both compounds [48]. In water, this compound can be hydrolyzed into gluconic acid, acidifying the environment and preventing bacterial growth [49–51]. The presence of this gene, encoding an enzyme capable of reversing this acidification – at least locally – may explain how *B. apis* is able to thrive in the presence of royal jelly [18, 52]. Alternatively, it is possible that *B. apis* is contributing to the production of gluconic acid in this environment. BLAST searches of the metatranscriptomes and metagenomes of bacteria in the “core” honey bee microbiome [3, 6] resulted in zero hits, indicating that none of the “core” microbiome members possess a homolog of this gene. The presence of gluconolactonase may help explain the unique distribution of *B. apis* within the hive. Another bee-associated unique gene is an HdeD family acid-resistance protein, which in *E. coli* participates in resistance to acids at high cell densities [53]. The presence of this gene in *B. apis* may indicate an adaptation to living in low pH environments – such as the queen bee digestive tract or royal jelly [54].

An AI-2 E family transporter was identified as unique to *Bombella* spp. AI-2 is an auto-inducer responsible for activating cascades associated with quorum sensing. While *B. apis* does not contain any AI-2 synthesis genes, the presence of an AI-2 E family transporter indicates that it may be responding to exogenous AI-2 produced by other bacteria, possibly in a competitive interaction. It is possible, also, that *B. apis* simply consumes this metabolite, using it as a source of carbon and energy. Bolstering the competition hypothesis is the presence of fusaric acid resistance (FUSC) genes in *B. apis*. Fusaric acid and its analogs can be quorum sensing inhibitors [55], so the presence of FUSC genes might be an adaptation that allows *B. apis* to evade quorum sensing inhibition attempts by other microbes. Alternatively, these FUSC genes may play a role in competition with fungal species. Fusaric acid is produced by several species of fungus and is antibacterial [56]. Therefore, the FUSC genes may play a role in *B. apis*’s protection of honey bee larvae and queens from infection with *Nosema* and other pathogens by allowing it to tolerate antibacterial capabilities of fungi and exert its antifungal properties [15, 19]. Interestingly, none of the canonical honey bee gut symbionts encode FUSC genes, further suggesting a unique role for this gene in *B. apis* among honey bee symbionts.

An invasion-associated locus B (Ialb) protein was identified as present in bee-associated AAB, but absent in *Saccharibacter spp*. In *Bartonella bacilliformis*, *ialb* mutants are impaired in their ability to colonize human erythrocytes, suggesting a role for this protein in eukaryotic cell invasion [57]. While it is not clear whether *Bombella* strains are ever intracellular, the presence of *ialb* suggests that it may have this capability.

The final set of genes of particular interest in this analysis is a complete Type I-E CRISPR/Cas cassette. To determine if this CRISPR/Cas cassette was active, we annotated the genomes for the presence of CRISPR arrays, and found that all of the genomes that have this CRISPR/Cas cassette contain multiple CRISPR arrays. It is possible that these CRISPR arrays were present in the most recent common ancestor of the *Bombella* clade and have simply remained in these current genomes; if that were the case, we would expect the spacers in these CRISPR arrays to be highly similar between all strains. However, if these arrays are part of an active CRISPR/Cas system, we would expect the spacers to differ from strain to strain, reflecting unique challenges encountered by each strain. To rule out the possibility that these arrays are ancestral, we aligned each spacer sequence from a given genome to all other spacer sequences from the other genomes and calculated the percent identity. The minimum best intergenomic match for any spacer was 40%, while the maximum was just 65% identical over the length of the spacer, indicating that the spacer sequences are unique from genome to genome and the CRISPR/Cas systems identified here are likely active and/or were incorporated after the *B. apis* strains diverged from one another. Importantly, the spacers also do not match the existing prophage genomes in the *Saccharibacter* or *Bombella* genomes and have homology to known, sequenced phages (Table S6). Top SEA-PHAGES hits for each spacer in each genome are quite distinct and reflect the fact that each of these sequenced *B. apis* strains has likely interacted with a different pool of phages.

*B. intestini* was isolated from a bumble bee gut, so we also looked at genes that were unique to this bacterium. There were a total of 65 genes that were unique to *B. intestini*, including a complete type IV secretion system (T4SS) and several genes involved in antibiotic production or resistance. Putative annotations of these 65 genes are in Table S3.

### Identification of horizontally transferred gene regions

Horizontal transfer of DNA between unrelated bacteria is a commonly known mechanism by which bacteria can acquire new traits and adapt to novel environments [58–61]. We identified two regions of phage origin, one in *S. floricola* and one in *B. apis* G7_7_3c (discussed above, Figure 2). To determine whether the bacteria in the *Bombella* clade have undergone other potential horizontal gene transfer (HGT) events, we determined the spatial distribution of genes of particular interest (e.g. clade-specific, species-specific, or strain-specific genes) across the bacterial genomes (Figure 3). Some of the genes specific to different clades occur in clusters, an indication that they may have originated elsewhere and been horizontally inherited as a chunk of contiguous DNA. We then looked for anomalies in sequence composition (%GC) and phylogeny to determine whether they were putatively horizontally transferred. Using this combination of methods, we identified a total of five HGT regions in the *Bombella* clade, which we have numbered 1-5 (see Table S4 for %GC and lineage probability index (LPI)-difference deviations for each gene in each HGT).

**Fig. 3.**
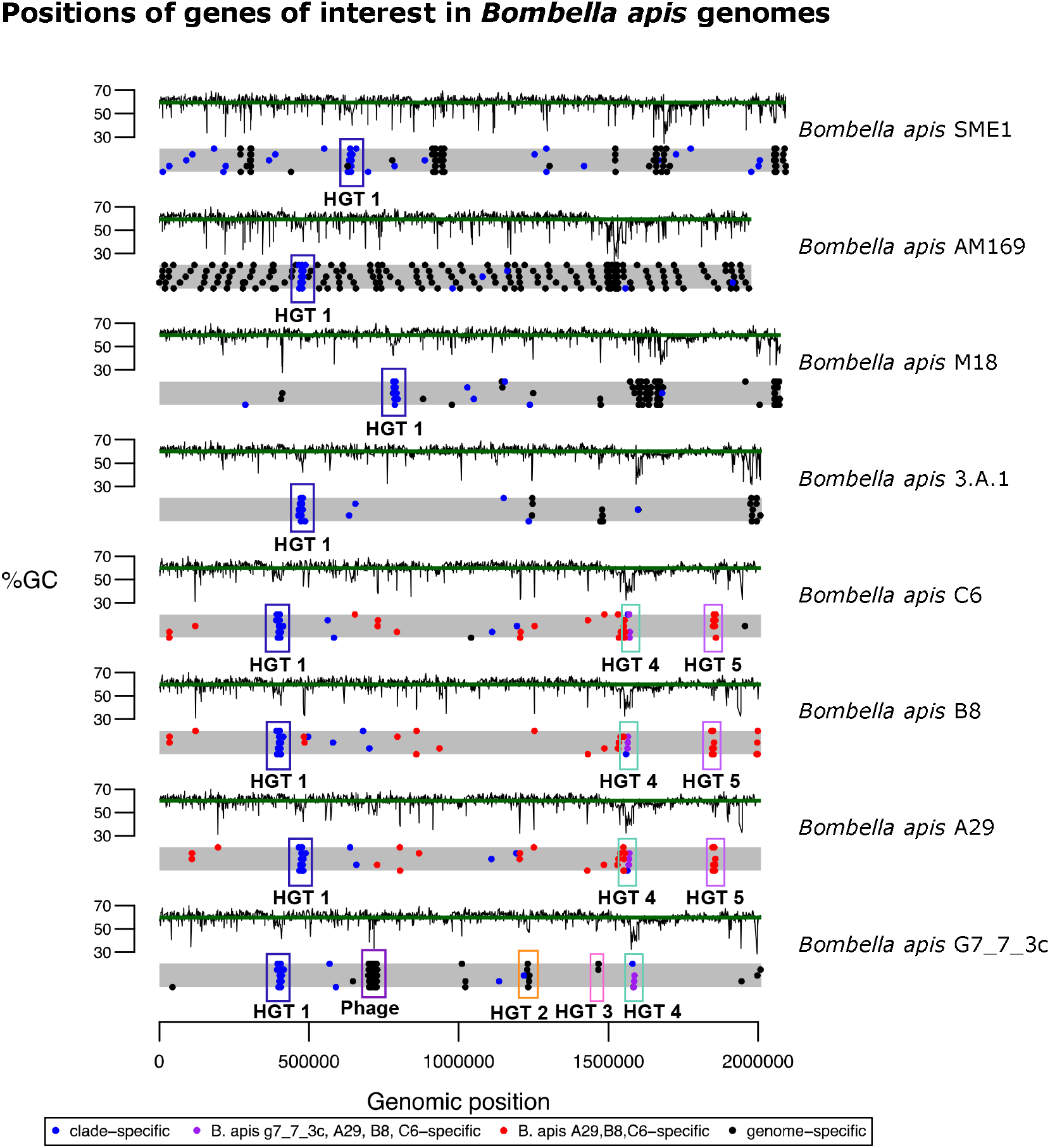
Genomic locations of genes of interest in *Bombella* and *Saccharibacter* genomes. Each gray bar is a representation of the genome, with each dot representing the location of a gene in each of four categories (see legend). Regions of interest mentioned in the text are highlighted and labeled. %GC for every gene is plotted above each genome representation, with the green line indicating the genome-wide average %GC.

HGT1 (Figure 4A) is present in all genomes in the *Bombella* clade, and contains 10 genes, although *B. apis* C6 is missing one of the genes (the second-to-last gene at the 3’ end of the HGT, annotated as an ABC transport auxiliary component). The three most 5’ genes show homology to YfaP (an uncharacterized conserved protein), SrfB (part of the surfactin antibiotic synthesis machinery), and an uncharacterized bacterial virulence factor. The genes in the 3’ half of this HGT contain a number of domains involved in membrane transport. We hypothesize that the two halves of this HGT work together to synthesize and export antibiotics as a form of defense or regulation of competing bacteria. Lending support to the hypothesis that this HGT is involved in defense or immunity is the fact that a CRISPR array lies immediately 5’ of this HGT in each genome (Table S5). Bacterial defense mechanisms tend to occur in clusters of “defense islands” [1] so the presence of this CRISPR array is perhaps a further indication of this HGTs role in bacterial immunity.

**Fig. 4.**
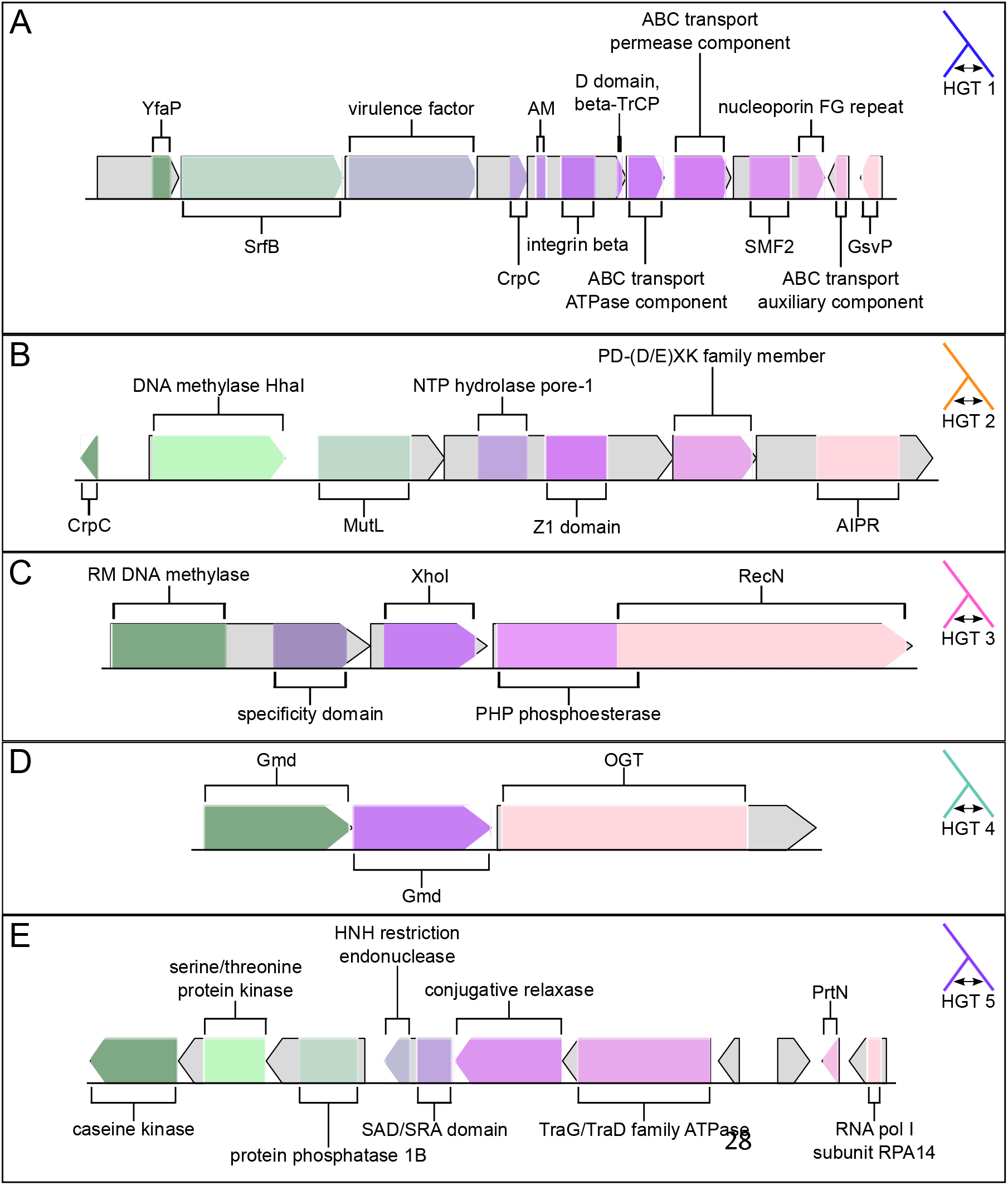
Gene models for each of 5 genomic regions of interest. Gene models are drawn to scale within each panel, but not across panels. A) HGT1. Abbreviations are: CrpC: cysteine rich protein C, AM: automated matches, SMF2: sulfatase modifying factor 2, GsvP: gas vesicle protein C. B) HGT2. Abbreviations are: CrpC: cysteine rich protein C, AIPR: abortive infection phage resistance protein. C) HGT3. Abbreviations are: RM: restriction-modification. D) HGT 4. Abbreviations are: Gmd: GDP-D-mannose dehydratase, OGT: O-linked N-acetylglucosamine transferase OGT. E) HGT5. Abbreviations are: SAD/SRA: SET and Ring finger Associated, PrtN: pyocin activator protein.

HGTs 2 and 3 (Figure 4B and 4C) are restricted solely to *B. apis* G7_7_3c and are both bacterial restriction-modification (R-M) systems. Bacterial R-M systems are a defense against invading DNA (i.e. bacteriophage). They act by methylating host DNA at specific sites; invading DNA with the same recognition site will be un-methylated, recognized as foreign, and targeted for degradation [62]. HGT2 contains 6 genes, which make up the core components of a bacterial (R-M) system. Interestingly, the domain architecture in this R-M system has been recognized as an evolutionary precursor to eukaryotic defenses against transposable elements [63]. HGT3 (Figure 4C) consists of 3 genes comprising 5 domains; the 5’-most gene consists of a predicted restriction-modification DNA methylase coupled to a specificity domain, the middle gene is predicted to be an XhoI restriction enzyme, and the 3’-most gene is a PHP phosphoesterase coupled to a RecN DNA repair ATPase. Taken together, it appears that HGTs 2 and 3 are responsible for recognition of and defense against foreign DNA.

HGT4 (Figure 4D) is present in *B. apis* strains G7_7_3c, A29, B8, and C6 and contains 3 genes: two GDP-D-mannose dehydratases (GMD) and an O-linked N-acetylglucosamine transferase (OGT). GMD plays a role in the metabolism of mannose and fructose, sugars commonly found in nectar [64]. The presence of GMD in *B. apis* genomes might allow for the consumption of nectar or nectar components by these bacteria. OGT, on the other hand, plays a role in post-translational modification of thousands of identified proteins [65]. However, while OGT-mediated post-translational modification is common in eukaryotes, it is far more rare in bacteria [66]. To date, only a handful of prokaryotic OGTs have been identified, and the targets of these OGTs remain unclear [67, 68]. Given the role OGTs play in eukaryotic post-translational modification and the fact that many bacterial effector proteins show homology to eukaryotic proteins [69], it is possible that the presence of OGT in *B. apis* represents a pathway for host-microbe interaction and symbiont-mediated protein modification.

HGT5 (Figure 4E) is unique to *B. apis* strains A29, B8, and C6, all strains that had been isolated from honey bee larvae. Like HGTs 1-3, HGT5 contains genes that may play a role in protection against foreign DNA. There are four genes in the 5’ section of HGT5, three of which are kinases, and the fourth contains a SAD/SRA domain in its 5’ end, and an HNH endonuclease domain in its 3’ end. In bacteria, the SAD/SRA domain is often found associated with an HNH domain [70] and it is thought that the two domains act together to recognize and cleave foreign DNA [63]. The 3’ section of HGT5 consists of a conjugative relaxase, a TraG/TraD family ATPase (a coupling protein involved in bacterial conjugation and/or T4SS), a homolog of the pyocin activator protein PrtN, a homolog of a yeast RNA polymerase I subunit, and two additional genes with no annotations. The presence of a PrtN homolog is particularly interesting, as in *Pseudomonas aeruginosa* pyocins are antibacterial agents, often acting to depolarize the membrane of target cells [71, 72]. Interestingly, one of the two unannotated genes in the 3’ region of HGT5 shows weak homology to a phage shock protein, which are proteins involved in the response to stress that may weaken the energy status of the cell [73]. This protein, then, may play a part in immunity to membrane depolarization. Given the presence in HGT5 of: an HNH endonuclease coupled to a SAD/SRA domain, a conjugative relaxase, a TraG/TraD family ATPase, a pyocin activator protein, and a protein with at least some homology to a phage shock protein, we hypothesize that it may play a role in pathogenesis or defense.

## Discussion

Here, we used the genomes of ten *Bombella* spp. and five *Saccharibacter* spp. to gain insight into the genomic changes associated with the transition to honey bee symbiosis in this group. We note several genomic differences – some of which were horizontally acquired – between bee-associated bacteria and the flower-associated *Saccharibacter* that may have allowed for the expansion of *B. apis* into previously unoccupied niches within the honey bee colony. These differences can be classified as changes that introduce: 1) novel metabolic capabilities, 2) defense and/or virulence mechanisms, and 3) mechanisms for interaction with other microbes and/or the host.

Metabolic genes identified here include gluconolactonase, which may allow for the de-acidification of royal jelly [48–51], and two copies of GMD, a gene that plays a role in the metabolism of mannose and fructose, components of nectar and honey [64]. Distinct defense and/or virulence mechanisms were identified, including: a CRISPR/Cas system, two R-M systems, and an HGT with some homology to known virulence mechanisms. Interestingly, the R-M systems were identified in the only strain in the clade that also contains a phage sequence (*B. apis* G7_7_3c). Restriction modification systems, like phages, can act as selfish genetic elements [74], so their presence in this genome may indicate that it was historically more permissive to invading DNA. These R-M systems may also have been coopted by the prophage to prevent super-infection with additional phages [75].

Genes involved in the interaction with other microbes and/or the host that we identified include: an AI-2 family transporter, fusaric acid resistance genes, *ialb*, and *ogt*. Given that *B. apis* does not encode any of the canonical genes for the production of quorum-sensing molecules, it seems likely that *B. apis* is responding to exogenous AI-2 (and/or fusaric acid and its analogs) produced by other members of the bee microbiome [76]. The *ialb* and *ogt* genes provide routes for interaction with the host, as *ialb* may play a role in eukaryotic cell invasion [57] and *ogt* is known to modify thousands of eukaryotic proteins [65]. Taken together, we hypothesize that the novel combination of metabolic, quorum-sensing, defense/virulence, and eukaryotic interaction genes in the *Bombella* clade genomes allowed for the utilization of a unique food source and protection from an onslaught of previously un-encountered challenges and facilitated the transition to honey bee association in this clade.

*Bombella apis* has been shown to benefit honey bee larval development and provide protection against *Nosema* and other pathogens [15, 19]. Some of the genes identified here, while allowing *B. apis* to transition to honey bee symbiosis, may also be related to its ability to protect the bee host from infection with *Nosema* or other pathogens. If indeed these genes are responsible for the transition to, and maintenance of, honey bee symbiosis, we would expect to see a modified evolutionary trajectory relative to those genes not involved in the symbiosis. We currently lack sufficient sampling of non-bee-associated bacteria in this clade to do such analyses; however, future studies addressing this question should allow for further elucidation of the genes involved in the transition to honey bee association. Those analyses, coupled with functional characterization of the genes of interest identified here, should lay the foundation for the development of beneficial intervention strategies in this economically critical insect.

## Supporting information

BLAST results for phage1

BLAST results for phage2

## Acknowledgements

We thank Amelia R.I. Lindsey for feedback on initial versions of this manuscript. E.A.S. was supported by a Faculty Research Support grant from Indiana University and by a USDA NIFA Postdoctoral Fellowship. This work was supported by NSF IOS 2005306 to I.L.G.N.

## Supplemental figures and table legends

**Table S1.** OrthoMCL clusters and gene counts for each type of COG in each genome

**Table S2.** Accession number, annotation source, annotation score, and putative annotation for each of the *Bombella*-specific genes identified

**Table S3.** Accession number, annotation source, annotation score, and putative annotation for each of the *Bombella intestini* genes identified

**Table S4.** %GC and LPI-difference standard deviations for each gene in each genome harbing each HGT.

**Table S5.** Positions and spacer counts for each CRISPR array identified in the *Bombella* clade genomes.

**Table S6.** SEA-PHAGES analysis of CRISPR spacers identified in each genome with top hit, hit length, and percent identity information.

**Table S7.** Top BLAST hits, percent identity, e-value, and query coverage, for each HGT identified in this analysis.

**Supplemental file S1.** BLAST results for genes present in phage 1, present in *Saccharibacter floricola.*

**Supplemental file S2.** BLAST results for genes present in phage 2, present in the *Bombella apis* G7_7_3c sequence.

## Notes

### Competing Interest Statement

The authors have declared no competing interest.

### Summary of Updates

In response to prior review we have substantially revised our manuscript in the following significant ways: We have included 6 additional genomes, expanding our outgroup Saccharibacter clade We have re-analyzed and re-run all analyses in this work using our additional genomes We have posted a bioRxiv preprint characterizing the taxonomic reassignment of this group We have run analyses of completeness and assembly for each genome We have looked for homology between our identified HGTs and existing databases We have identified possible phage from which our CRISPR spacers are derived

